# Catalase Activity is Critical for *Proteus mirabilis* Biofilm Development, EPS Composition, and Dissemination During Catheter-Associated Urinary Tract Infection

**DOI:** 10.1101/2021.03.22.436542

**Authors:** Ashley N. White, Brian S. Learman, Aimee L. Brauer, Chelsie E. Armbruster

**Author notes:** Correspondence: Chelsie Elizabeth Armbruster, 955 Main Street, Room 5218, Buffalo, NY 14203.

## Abstract

*Proteus mirabilis* is a leading uropathogen of catheter-associated urinary tract infections (CAUTIs), which are among the most common healthcare-associated infections worldwide. A key factor that contributes to *P. mirabilis* pathogenesis and persistence during CAUTI is the formation of catheter biofilms, which provide increased resistance to antibiotic treatment and host defense mechanisms. Another factor that is important for bacterial persistence during CAUTI is the ability to resist reactive oxygen species (ROS), such as through the action of the catalase enzyme. Potent catalase activity is one of the defining biochemical characteristics of *P. mirabilis,* and its single catalase gene (*katA*) was recently identified as a candidate fitness factor for UTI, CAUTI, and bacteremia. Here we show that disruption of *katA* results in increased ROS levels, increased sensitivity to peroxide, and decreased biofilm biomass. The biomass defect was due to a decrease in extracellular polymeric substances (EPS) production by the Δ*katA* mutant, and specifically due to reduced carbohydrate content. Importantly, the biofilm defect resulted in decreased antibiotic resistance *in vitro* and a colonization defect during experimental CAUTI. The Δ*katA* mutant also exhibited decreased fitness in a bacteremia model, supporting a dual role for catalase in *P. mirabilis* biofilm development and immune evasion.

## Introduction

*Proteus mirabilis* is a Gram-negative bacterium that is present in numerous environments such as soil, water, and the intestinal tracts of both humans and animals (1). However, *P. mirabilis* is also a leading uropathogen of catheter-associated urinary tract infections (CAUTIs), which are one the most common healthcare-associated infections worldwide (2–4). CAUTIs account for up to 80% of all nosocomial UTIs, and *P. mirabilis* CAUTIs are often complicated by bladder and kidney stone formation (urolithiasis), permanent renal damage, and progression to life-threatening bacteremia and sepsis (5–8). For instance, *P. mirabilis* is responsible for 12-31% of bacteremias in nursing home residents and is associated with a one-year mortality rate ranging from 10 to 66% (9–13).

*P. mirabilis* CAUTIs have proven difficult to treat in part due to their increasing antibiotic resistance, as well as resistant biofilm formation on the catheter surface (14–17)*. P. mirabilis* is particularly well adapted for biofilm formation, causing the encrustation and blockage of urethral catheters by the formation of crystalline biofilm structures (6, 18–22). Notably, biofilms containing *P. mirabilis* have been detected on catheters even after antibiotic treatment (23–25). As *P. mirabilis* biofilms play a significant role in both the pathogenesis and treatment of CAUTI (4, 6, 26, 27), it is critical to investigate factors that influence biofilm development and antibiotic resistance in this species.

Biofilm formation is a multistep process, involving irreversible microbial attachment to a substrate, development of microcolonies, the production of extracellular polymeric substances (EPS), maturation, and dispersal, which is thought to contribute to persistent infection during CAUTI and dissemination to the bloodstream (6, 28). The EPS matrix is highly hydrated (98% water) and largely comprised of proteins, carbohydrates, and extracellular DNA (eDNA); however, composition varies depending on bacterial strain, environmental conditions, and maturation stage of the biofilm (29). EPS is considered to be a critical component of bacterial biofilms, as it defines the biofilm structure and is essential for the overall integrity and function of the biofilm (19, 28, 30).

The EPS matrix also plays a substantial role in the increased antimicrobial resistance exhibited by bacterial biofilms (17). There are several ways in which biofilm formation impacts antimicrobial susceptibility. First, the antimicrobial agents must diffuse through the EPS matrix in order to act on the organisms within the biofilm. The EPS effectively acts as a shield that prevents diffusion of antimicrobials either by binding and chemically inhibiting the antimicrobial molecules or by limiting their rate of infiltration (31, 32). The biofilm-associated bacteria also experience reduced growth rates and are less metabolically active, which limits the efficacy of many antimicrobial agents (33). Lastly, the environment that immediately surrounds the biofilm may provide conditions that further protect the biofilm-associated bacteria (33). As a result, biofilm formation on indwelling medical devices poses a serious challenge due to the increased resistance of biofilm-associated organisms to antimicrobial agents (28).

Another factor that contributes to bacterial persistence and pathogenicity is the ability to resist reactive oxygen species (ROS) (34). For instance, many bacterial species produce a catalase enzyme, which reduces hydrogen peroxide (H_2_O_2_) to water (H_2_O) and oxygen (O_2_), and has been shown to protect biofilm associated *Pseudomonas aeruginosa* by preventing full penetration of H_2_O_2_ through the biofilm matrix (34). Hydrogen peroxide is the longest-lived reactive oxygen species, and it can cause critical damage to both the bacterial cell membrane and essential cellular macromolecules such as proteins, lipids, and nucleic acids (35, 36). ROS can also contribute to the effectiveness of antibiotic mediated killing. For example, antibiotic exposure induces oxidative stress in *P. aeruginosa,* and antibiotic susceptibility was increased by disruption of a gene involved in reducing H_2_O_2_ (*ahpC*), indicating that ROS generation contributed to drug lethality (37). It has therefore been proposed that interfering with bacterial protection against ROS could improve antibiotic treatment, even for biofilms (37–40).

Potent catalase activity is one of the defining biochemical characteristics of *P. mirabilis* (41), and there are several indications that the single catalase gene (*katA*) of *P. mirabilis* is important for both fitness and pathogenicity during infection. For instance, *katA* was found to be upregulated during infection in a mouse model of UTI (42), it was identified as a candidate fitness factor during both single-species and polymicrobial CAUTI (43), and it was detected as an infection-specific fitness factor for survival in the bloodstream (44). Despite this evidence, the exact role of catalase in *P. mirabilis* pathogenesis had yet to be experimentally validated. In this study, we disrupted the *katA* gene in *P. mirabilis* strain HI4320 and investigated the contribution of catalase to bacterial growth, motility, ROS tolerance, biofilm formation, antibiotic resistance, pathogenicity within the urinary tract, and fitness within the bloodstream. We show that loss of *P. mirabilis* catalase activity does not alter *in vitro* growth or planktonic antibiotic susceptibility, but peroxide detoxification is critical for production of a mature biofilm. The absence of catalase severely alters the composition of the EPS matrix, resulting in increased antibiotic susceptibility of biofilm-associated bacteria. Additionally, we demonstrate that catalase contributes to the pathogenesis of *P. mirabilis* during CAUTI and fitness during bacteremia.

## Results

### Generation and characterization of a P. mirabilis ΔkatA mutant

In order to investigate the importance of catalase activity to *P. mirabilis* growth, biofilm formation, and pathogenesis, a mutant was generated in *P. mirabilis* strain HI4320 to disrupt the catalase gene (*katA*) by insertion of a kanamycin resistance cassette. Disruption of the *katA* gene and resulting catalase activity were confirmed by PCR and a catalase foam height assay (Fig. 1A). As expected, the Δ*katA* mutant was catalase-negative when exposed to hydrogen peroxide, and providing the *katA* gene on a plasmid restored catalase activity. The complemented activity was more potent than that of the parental strain, which is likely due to plasmid copy number. The Δ*katA* mutant grew similarly to wild-type *P. mirabilis* in LB broth and did not exhibit any defects in swimming motility, swarming motility, or urease activity (Fig. 1B-E), indicating that disruption of the *katA* gene does not affect standard laboratory behaviors of *P. mirabilis*. Disruption of *katA* also had no impact on expression of either adjacent gene by qRT-PCR (Fig. S1). Taken together, these data indicate that targeted disruption of *katA* abrogates catalase activity without obvious polar effects.

**Figure 1:**
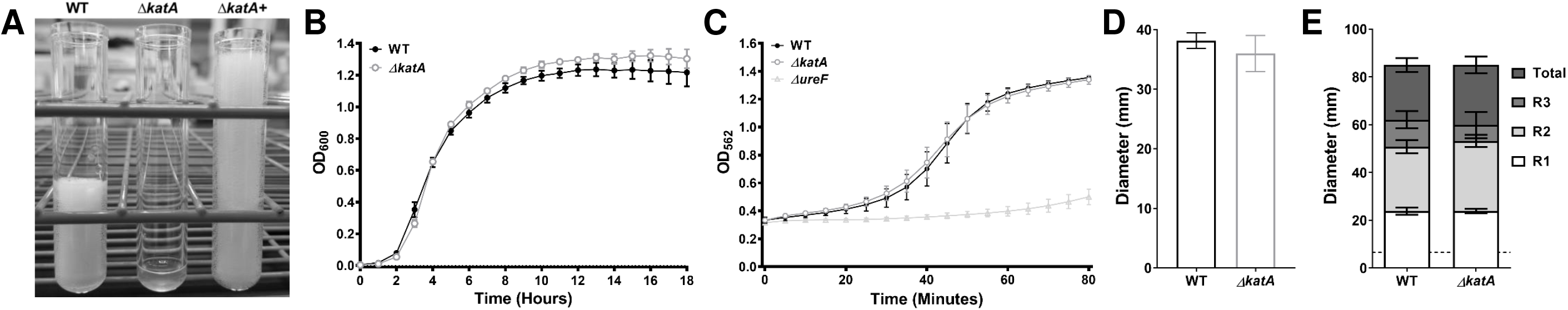
Characterization of a P. mirabilis ΔkatA mutant. The Δ*katA* mutant was assessed for catalase activity (A), growth in LB (B), urease activity (C), swimming motility (D), and swarming motility (E) in comparison to wild-type *P. mirabilis*. (A) An SDS/peroxide foam height assay was conducted to provide phenotypic validation of successful generation of a *P. mirabilis ΔkatA* mutant and plasmid generated Δ*katA+* complemented strain. (B) Bacterial growth in LB at 37°C was assessed by measurement of OD_600_ at hourly intervals for 18 hours. Error bars represent mean ± standard deviation (SD) of 6 independent experiments with 6 technical replicates each (representative graph shown). No significant differences in growth were determined by two-way ANOVA with Dunnett’s test for multiple comparisons. (C) Urease activity in human urine was assessed by measurement of phenol red color change (OD_562_) at 5-minute intervals for 80 minutes, and a *P. mirabilis ΔureF* mutant was included as negative control. Error bars represent mean ± SD of 2 independent experiments with 6 technical replicates each. No significant differences between WT and Δ*katA* were identified by two-way ANOVA. (D and E) Bacteria were cultured in LB broth overnight and stab inoculated into motility agar (D) or inoculated onto the surface of swarm agar plates (E). Motility diameter was measured in millimeters after 16 hours of incubation at 30°C (D) or 37°C (E). R1-3 indicate individual swarm ring diameters. Error bars represent mean ± SD for 3 independent experiments with at least 2 replicates each. No significant differences in motility diameter were determined by Student’s t test (D) or two-way ANOVA (E).

### Disrupting katA increases sensitivity to hydrogen peroxide and steady-state ROS levels

In order to define the limits of hydrogen peroxide each strain could withstand, we conducted a hydrogen peroxide sensitivity growth curve wherein each strain was cultured in LB broth with increasing concentrations of H_2_O_2_. The Δ*katA* mutant was susceptible to ∼2 mM H_2_O_2_ and completely inhibited by ∼3 mM (Fig. 2B, C). In contrast, ∼109 mM H_2_O_2_ was needed to impair growth of wild-type *P. mirabilis* and ∼326 mM H_2_O_2_ for complete growth inhibition (Fig. 2D, E). Importantly, providing the *katA* gene on a plasmid allowed the Δ*katA* mutant to tolerate higher H_2_O_2_ concentrations than the wild-type strain (Fig. 2E), again likely due to a high plasmid copy number.

**Figure 2:**
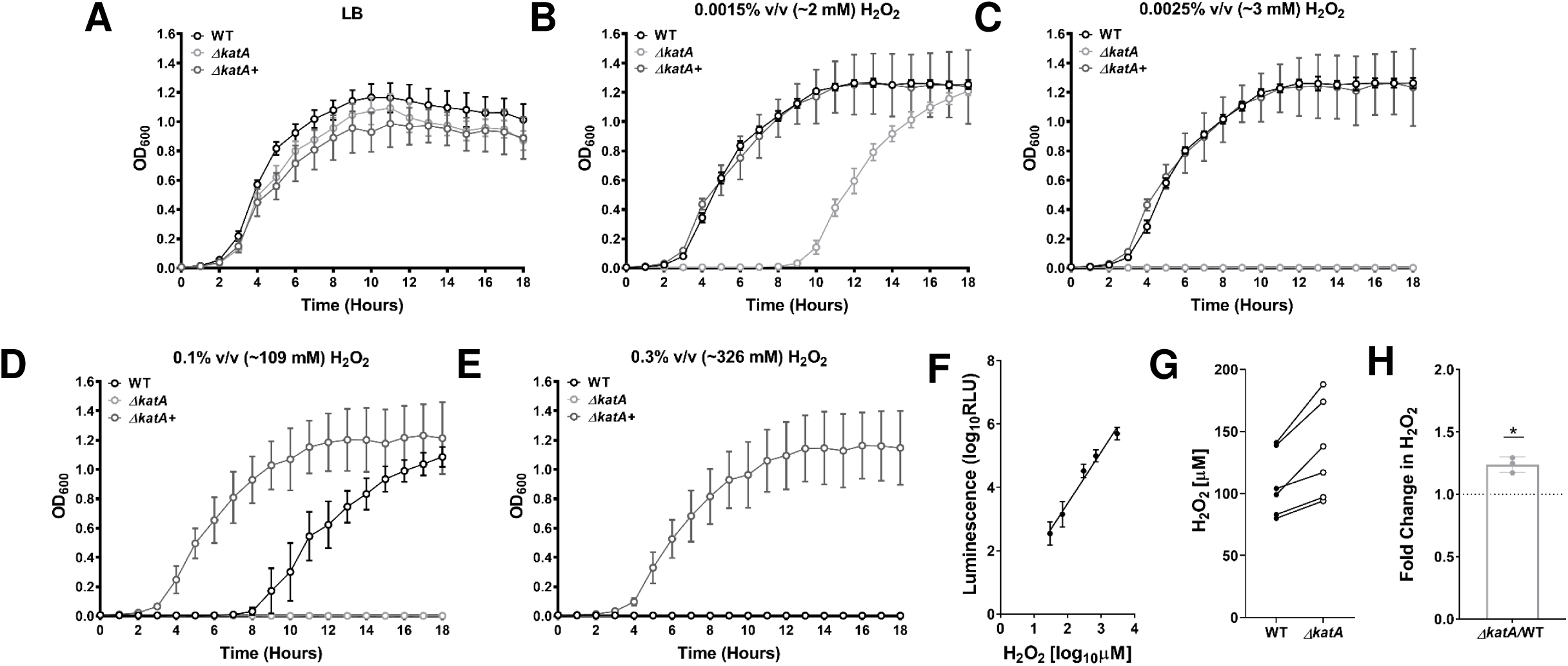
Disruption of katA increases sensitivity to hydrogen peroxide and steady-state ROS levels. (A-E) Growth at 37°C was assessed by measurement of OD_600_ at hourly intervals for 18 hours for wild-type (WT, black), Δ*katA* mutant (light gray), and plasmid generated Δ*katA+* complemented strain (dark gray) in (A) plain LB or in the presence of increasing concentrations of hydrogen peroxide (H_2_O_2_) as follows: (B) 0.0015% v/v (∼2 mM), (C) 0.0025% v/v (∼3 mM), (D) 0.1% v/v (∼109 mM), and (E) 0.3% v/v (∼326 mM). Error bars represent mean ± standard deviation (SD) of 3 independent experiments with 6 technical replicates each. (F) Peroxide standard curve for measurement of reactive oxygen species in bacterial cultures. (G) H_2_O_2_ concentrations present in cultures of WT (dark gray) and Δ*katA* (white) after a 4-hour incubation in LB broth. Individual data points represent 3 independent experiments with technical duplicates, and the WT and Δ*katA* values from each experiment are connected with a black line. (H) Average fold change in Δ*katA* H_2_O_2_ levels compared to WT. * P<0.05 by one-sample t test.

A luminescence-based H_2_O_2_ assay was next used to determine the extent of H_2_O_2_ accumulation in wild-type *P. mirabilis* and the Δ*katA* mutant during growth in broth culture to gauge the contribution of catalase activity to detoxification of endogenously-produced ROS. For these experiments, a standard curve was generated by supplementing Δ*katA* cultures with increasing concentrations of peroxide at the time of ROS measurement (Fig. 2F). After 4 hours of culture in LB broth, samples from wild-type *P. mirabilis* contained an average of 109 ± 26.9 µM H_2_O_2_, while the Δ*katA* samples contained an average of 135 ± 39.7 µM H_2_O_2_ (1.24 fold greater than wild-type, Fig. 2G, H). Notably, the concentration of H_2_O_2_ in the Δ*katA* samples is ∼15-fold lower than the concentration at which growth of Δ*katA* was perturbed (∼2mM, as seen in Fig.2B), suggesting that the loss of catalase activity likely does not induce a significant state of oxidative stress during growth in LB broth. However, it could cause the Δ*katA* mutant to be more susceptible to additional ROS insult or other stressors.

### The ΔkatA mutant exhibits a defect in biofilm development

As biofilm formation is a critical component of *P. mirabilis* pathogenesis during CAUTI, we next assessed biofilm formation by the Δ*katA* mutant at 2, 4, 6, 12, and 20 hours (Fig 3). By crystal violet staining, wild-type *P. mirabilis* established a robust biofilm over the 20 hour incubation in LB broth, but the Δ*katA* mutant exhibited a significant decrease in total biomass that was detectable at 4 hours and remained evident for the rest of the time course (Fig. 3A). Importantly, assessment of viable bacteria within the biofilms at each time point revealed that the defect was not due to a decrease in bacterial CFUs, and therefore likely represents a defect in biofilm architecture (Fig. S2). These results were also recapitulated for biofilms established in pooled human urine, as a physiologically-relevant growth medium for CAUTI (Fig. 3B).

**Figure 3:**
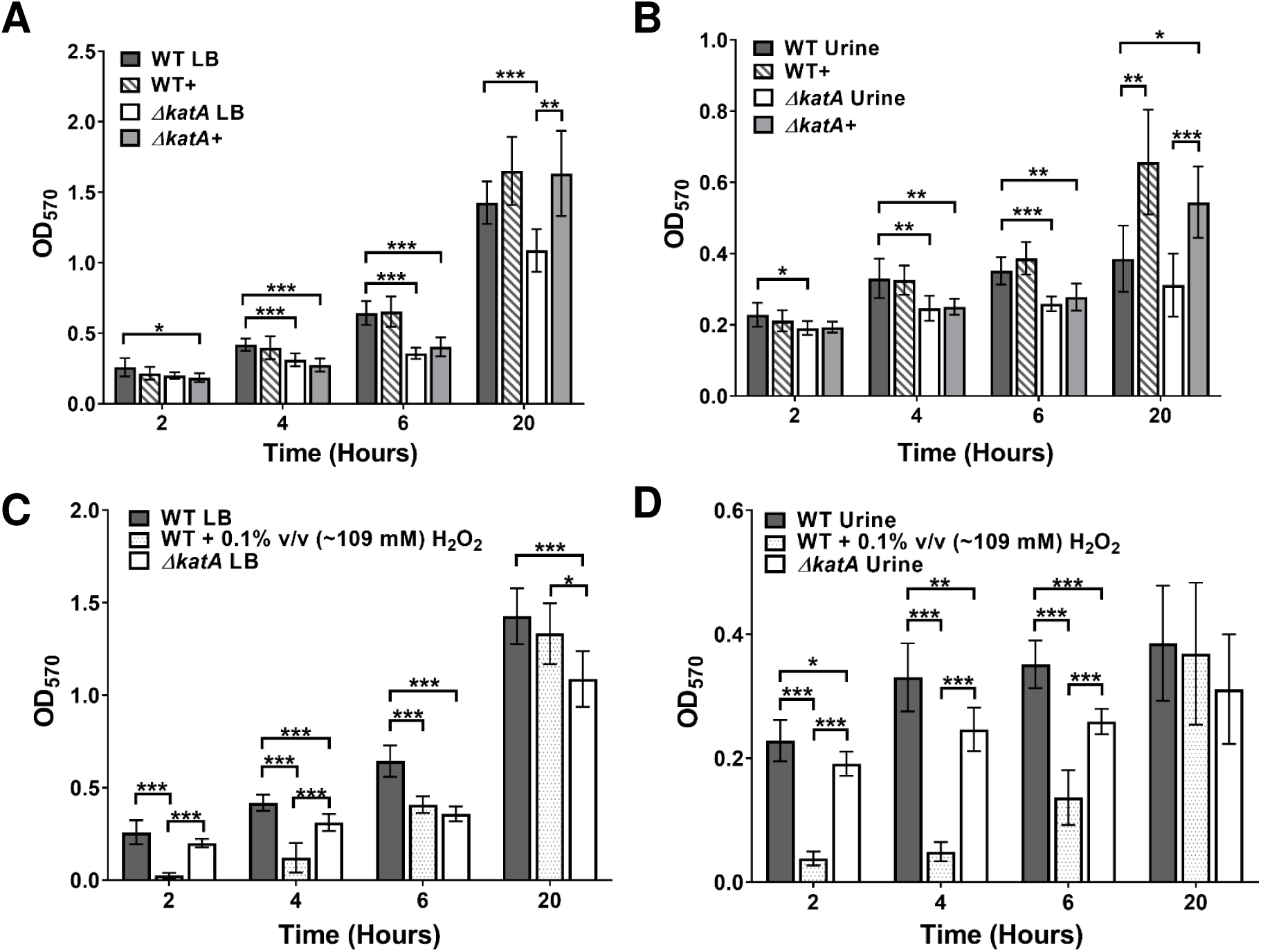
ΔkatA exhibits a defect in biofilm development. (A,B) Biofilms of WT (gray), WT+ (+bovine catalase, gray stripes), Δ*katA* (white), and Δ*katA+* (+ bovine catalase, light gray) were formed over a timecourse of 20 h in (A) LB or (B) pooled human urine. Biofilm biomass was assessed at 2, 4, 6, 12, and 20 h via crystal violet staining and detected by OD_570_. (C, D) Biofilms of WT (gray), WT + 0.1% v/v (∼109 mM) H_2_O_2_ (gray dots), and Δ*katA* (white) were formed over a timecourse of 20 h in (C) LB or (D) pooled human urine. (A-D) Error bars represent mean and SD of 3 independent experiments with 3 replicates each. *P<0.05, ** P<0.01, ***P<.001 by two-way ANOVA with Sidak’s test for multiple comparisons.

To define the specific contribution of peroxide detoxification to *P. mirabilis* biofilm formation, wild-type *P. mirabilis* and the Δ*katA* mutant were supplemented with an amount of bovine catalase that results in the equivalent foam height produced by 1×10^8^ CFU/ml of wild-type *P. mirabilis* (∼0.7 mg, ∼40,000 units). The addition of bovine catalase had no impact on biofilm development by wild-type *P. mirabilis* in LB, but fully restored biofilm biomass for the Δ*katA* mutant in both LB and human urine, albeit not until the 20 hour time point (Fig. 3A, B). These data suggest that catalase is important for later stages of biofilm development and maturation rather than the initial stages of attachment and microcolony formation. Importantly, the contribution of bovine catalase was confirmed to specifically derive from enzymatic activity, as both 10 kDa filtration and heat inactivation of the catalase solution abrogated complementation of the Δ*katA* mutant biofilm defect (Fig. S3).

To verify that the contribution of catalase to *P. mirabilis* biofilm development pertains specifically to peroxide detoxification, we next sought to determine if the addition of a non-lethal concentration of peroxide (∼109 mM, as seen in Fig. 2D) could impair biofilm formation by wild-type *P. mirabilis*. The addition of peroxide at the time of inoculation significantly decreased the *P. mirabilis* biofilm biomass for the first 6 hours of biofilm development, after which time enough of the peroxide was likely broken down to restore full biofilm development (Fig. 3C). Similar results were also recapitulated in pooled human urine (Fig. 3D). Taken together, these data confirm that the biofilm defect of the Δ*katA* mutant is specifically due to the loss of catalase activity and peroxide detoxification, and likely pertains to biofilm maturation.

We next sought to determine if the biofilm defect of the Δ*katA* mutant may be the result of increased oxidative stress. Throughout the time course of biofilm development, samples were taken from the Δ*katA* mutant and wild-type *P. mirabilis* to examine the relative expression level of two additional peroxide detoxifying enzymes (alkylhydroperoxidases *ahpC* PMI1213 and PMI0073) and three superoxide dismutases (*sodA*, *sodB*, *sodC)*. Expression of each gene was normalized to *rpoA,* and relative expression over time was determined for the wild-type strain alone and the Δ*katA* mutant in comparison to the wild-type (Fig S4). Expression of genes involved in peroxide detoxification (*katA*, PMI1213 and PMI0073) remained unaffected in the wild-type strain over the time course of biofilm development (Fig. S4A-C); however, expression of genes involved in superoxide detoxification (*sodA*, *sodB*, and *sodC*) were significantly upregulated over time (Fig. S4D-F). This suggests that wild-type *P. mirabilis* experiences some degree of oxidative stress during biofilm formation, and is primarily due to superoxide. If disruption of catalase and resulting accumulation of endogenous peroxide generated substantial oxidative stress, we would expect to see increased expression of some or all of these genes in the Δ*katA* mutant relative to wild-type *P. mirabilis*. However, none of the selected genes exhibited a significantly different expression pattern in Δ*katA* biofilms in comparison to the wild-type biofilms (Fig. S4H-L). Thus, while *P. mirabilis* does exhibit signs of oxidative stress during biofilm formation, loss of catalase activity does not cause a substantial increase in oxidative stress.

### ΔkatA exhibits a defect in biofilm EPS composition

Since the Δ*katA* mutant exhibited a defect in biofilm biomass independent of cell viability or oxidative stress, these data led us to investigate the overall composition of the biofilm. Aside from bacterial aggregates, the other major component of bacterial biofilms is the matrix of extracellular polymeric substances (EPS) that coats and protects the bacterial cells from the surrounding environment. To qualitatively assess biofilm composition, we utilized scanning electron microscopy (SEM). Biofilms formed by wild-type *P. mirabilis* for 20 hours were heavily coated with EPS (Fig. 4A), while biofilms formed by the Δ*katA* mutant displayed a similar degree of aggregated clusters of bacteria but were lacking the extensive EPS coating of the wild-type biofilms (Fig. 4B). Importantly, the addition of bovine catalase at the time of inoculation slightly enhanced EPS production by wild-type *P. mirabilis* (Fig 4C) and fully restored EPS production by the Δ*katA* mutant (Fig 4D). These data indicate that the biomass defect of the Δ*katA* mutant is likely due to a decrease in the production of EPS.

**Figure 4:**
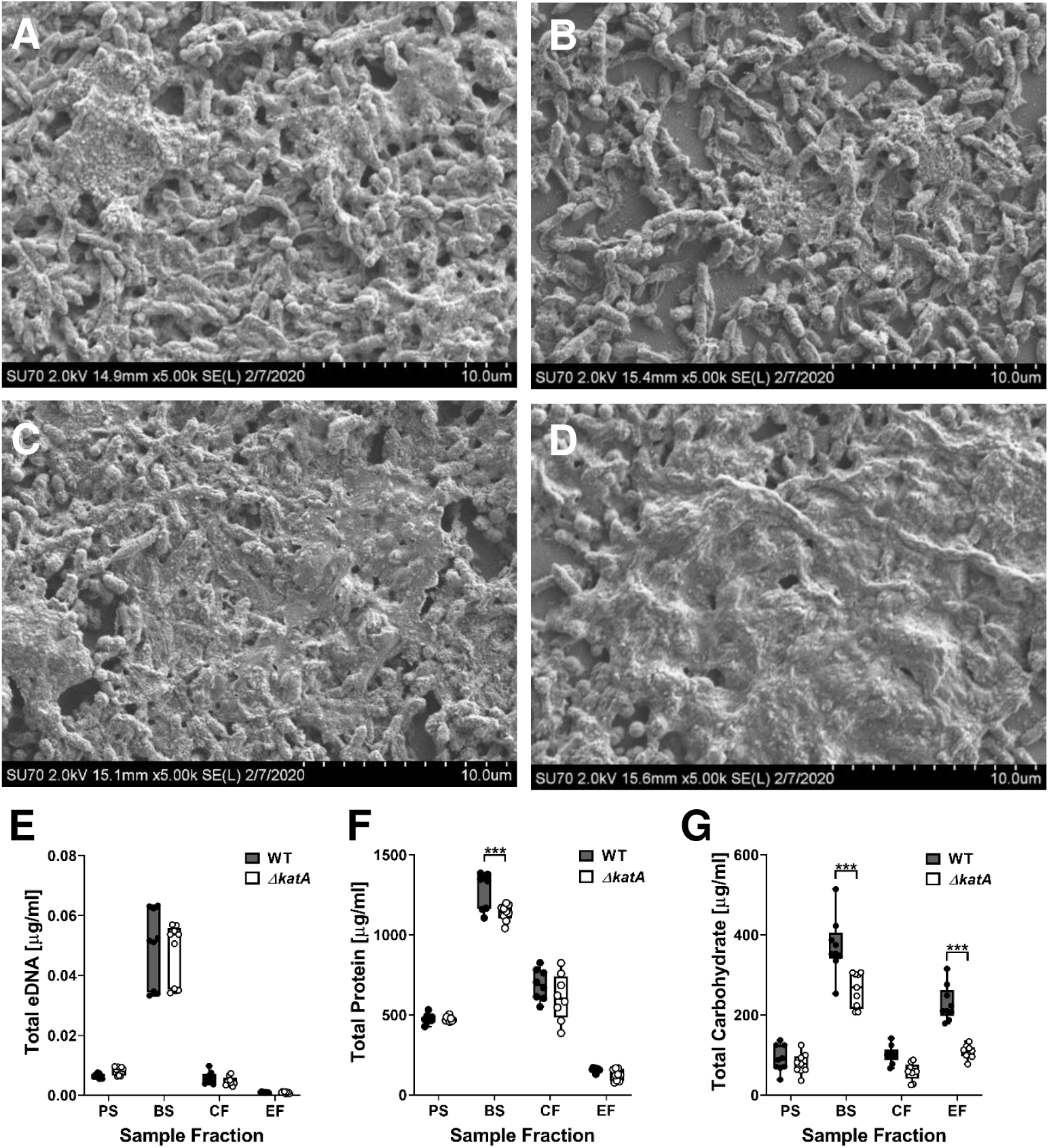
ΔkatA exhibits a defect in biofilm EPS production. (A-D) Representative SEM images of 20 h biofilms formed by (A) WT, (B) Δ*katA*, (C) WT+ bovine catalase, and (D) Δ*katA+* bovine catalase. (E-G) WT (black circles) and Δ*katA* (white circles) were incubated for 20 h with aeration or stationary to generate planktonic suspensions (PS) and biofilm (BS) suspensions. EPS was further extracted from the biofilm suspension to generate a cellular fraction (CF) and EPS fraction (EF) for measurement of eDNA (E), protein (F), and carbohydrate (G). Box and whisker plots display the combined data from 3 independent experiments with 3 technical replicates each. ***P<.001 by two-way ANOVA with Sidak’s test for multiple comparisons.

The exact composition of *P. mirabilis* HI4320 biofilm EPS has not been fully elucidated in the literature; however, the EPS of bacterial biofilms is generally comprised of proteins, carbohydrates, and extracellular DNA (eDNA). To assess biofilm EPS composition, wild-type *P. mirabilis* and the Δ*katA* mutant were incubated for 20 hours in LB broth with aeration (planktonic suspension, PS) or stationary in wells of a 24-well plate from which a total biofilm sample was collected (biofilm suspension, BS). The remaining biofilm material was then treated with formaldehyde to fix the bacterial cells and prevent lysis during subsequent steps of EPS extraction, and NaOH to promote dissociation of the EPS from the biofilm and increase its solubility. Following these treatments, a cellular fraction (CF) and dialyzed EPS fraction (EF) were collected. No differences in eDNA concentration were observed between strains in any of the tested fractions (Fig. 4E), and total protein measurement revealed only a slight decrease in the Δ*katA* biofilm suspension compared to wild-type (Fig. 4F). However, total carbohydrate measurement revealed a substantial decrease in the biofilm suspension and EPS fraction of the Δ*katA* mutant compared to wild-type *P. mirabilis* (Fig. 4G). Thus, the biofilm biomass defect of the Δ*katA* mutant derives from a decrease in EPS carbohydrate content.

### ΔkatA biofilms exhibit increased antibiotic susceptibility

Due to the enhanced antimicrobial resistance of bacteria in a biofilm mode of growth, we sought to determine the impact of the Δ*katA* biofilm biomass defect on antibiotic susceptibility. Wild-type *P. mirabilis* and the Δ*katA* mutant were treated with two antibiotics that target the bacterial cell wall (ampicillin and ceftriaxone) and one that targets DNA replication (ciprofloxacin) during planktonic growth or after 20 hours of biofilm growth. The Δ*katA* mutant did not exhibit substantial differences in susceptibility to ampicillin or ciprofloxacin during planktonic growth, although a slight decrease was detected for ceftriaxone at 0.02 and 0.01 µg/ml (Fig. 5A-C). However, Δ*katA* 20 h biofilms were more susceptible to all three antibiotics, suggesting that the decrease in biofilm biomass impacts antibiotic susceptibility (Fig. 5D-F). To further explore the importance of the biofilm defect for antibiotic susceptibility, biofilms formed by wild-type and the Δ*katA* mutant were also treated after only 4 hours of development, as this was the earliest time point corresponding to a significant decrease in biofilm biomass for the Δ*katA* mutant. Earlier exposure to antibiotics resulted in greater killing of both strains and further magnified the defect of the Δ*katA* mutant (Fig. 5G-I). Importantly, the addition of bovine catalase restored the antibiotic resistance of the Δ*katA* mutant biofilms to an equivalent or even greater level than the wild-type strain (Fig. 5G-I). Thus, we conclude that loss of *katA* results in a biofilm-specific increase in antibiotic susceptibility due to disruption of EPS production and biofilm maturation.

**Figure 5:**
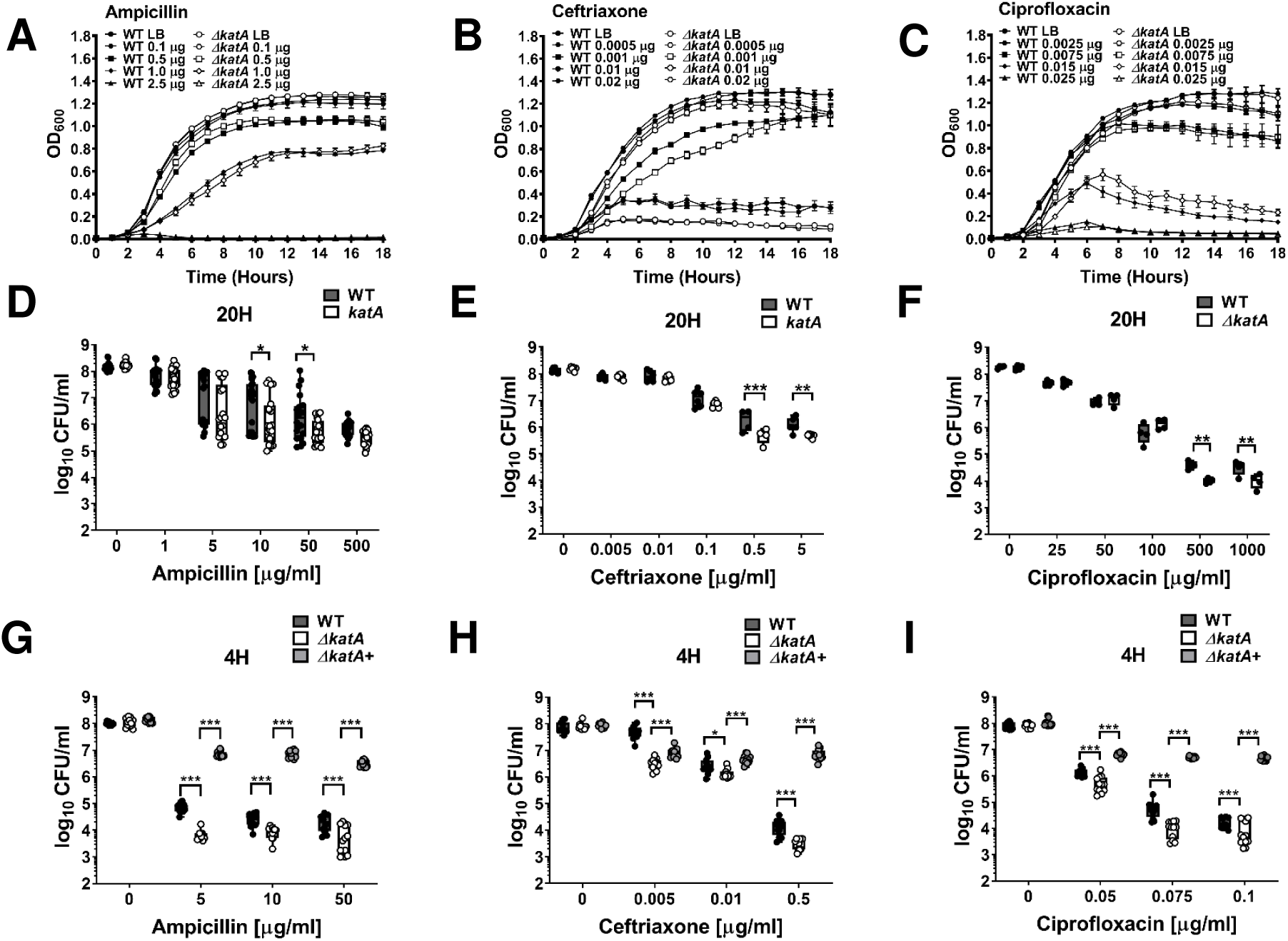
Δ*katA* biofilms exhibit increased antibiotic susceptibility. (A-C) Growth at 37°C was assessed by measurement of OD_600_ at hourly intervals for 18 hours for WT (black symbols) and Δ*katA* (white symbols) in the presence of increasing concentrations (µg/ml) of (A) ampicillin, (B) ceftriaxone, or (C) ciprofloxacin. Error bars represent mean ± standard deviation (SD) for 6 technical replicates, and graphs are representative of at least 3 independent experiments. Data were analyzed by two-way ANOVA with Tukey’s test for multiple comparisons. (D-I) Biofilms were established by WT and Δ*katA* for 20 h (D-F) or 4 h (G-I), followed by treatment with increasing concentrations of ampicillin (D and G), ceftriaxone (E and H), or ciprofloxacin (F and I) for an additional 20 h period. Each black (WT), white (Δ*katA*), and light gray (katA+ bovine catalase) circle represents the CFU/ml recovered from an individual biofilm, and box and whisker plots display combined results from at least 3 independent experiments with at least 3 technical replicates each. * P<0.05, ** P<0.01, ***P<.001 by two-way ANOVA with Sidak’s test for multiple comparisons.

### The biofilm biomass defect of the ΔkatA mutant can be recapitulated on Foley catheters

The bacterial biofilms that drive persistent infection in CAUTI are most often formed on the surface and within the lumen of the catheters themselves. We therefore sought to determine if the Δ*katA* biofilm biomass defect is similarly present when biofilms are formed directly on a silicone Foley catheter. Sterile Foley catheters (size 14 French) were cut into 15-mm segments, suspended in the wells of a 24 well plate, and inoculated with either wild-type *P. mirabilis* or the Δ*katA* mutant in LB broth. Biofilms were established for 20 hours, after which time the catheter segments were removed, air dried, and biofilm biomass was measured using an optimized crystal violet assay. Wild-type *P. mirabilis* established robust biofilms on the silicone catheter segments, and the Δ*katA* mutant again exhibited a substantial biomass defect that could be complemented by the addition of bovine catalase at the time of inoculation (Fig. 6A, B). Thus, the Δ*katA* biofilm biomass defect is present on a physiologically-relevant biofilm substrate for CAUTI.

**Figure 6:**
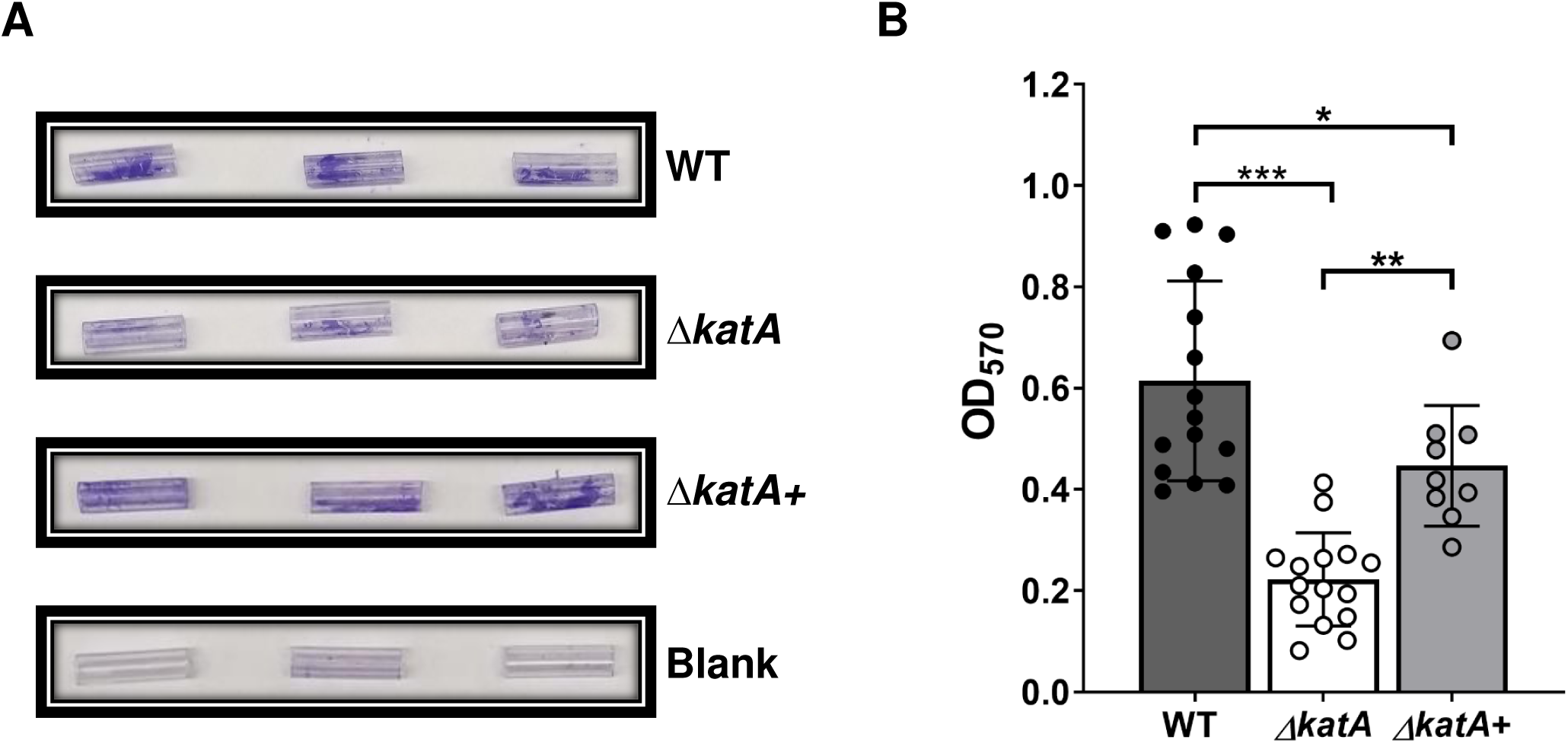
ΔkatA exhibits a biofilm biomass defect on silicone catheters. (A) Representative images of crystal violet-stained 15 mm silicone Foley catheter segments after 20 h incubation with WT, Δ*katA*, Δ*katA+* (bovine catalase), or plain LB (blank). (B) Biomass of 20 h WT (gray), Δ*katA* (white), ΔkatA+ (bovine catalase, light gray) catheter biofilms as assessed by crystal violet staining at OD_570_. Error bars represent mean and standard deviation (SD) of at least 3 independent experiments with at least 3 technical replicates each. ** P<0.01, ***P<.001 by one-way ANOVA with Tukey’s test for multiple comparisons.

### Catalase contributes to both pathogenesis and fitness in vivo

Considering the important contribution of bacterial biofilms to CAUTI in humans, we next sought to determine the contribution of *P. mirabilis* catalase to colonization and pathogenicity during experimental CAUTI. To test this, we utilized a mouse model of CAUTI as previously described (45). Briefly, female CBA/J mice were transurethrally inoculated with 1 x 10^5^ CFU of either wild-type *P. mirabilis* or the Δ*katA* mutant, and a 4-mm segment of sterile silicone catheter tubing was inserted into the bladder at the time of inoculation to recapitulate CAUTI. Weight loss was monitored daily (Fig. S5) and mice were euthanized at 24 or 96 hours post inoculation (hpi) to quantify bacterial burden in urine, bladder, and kidneys, as well as the spleen as an indication of bacteremia. Infection with wild-type *P. mirabilis* resulted in slightly greater weight loss at 24 hpi than infection with the Δ*katA* mutant, but no significant differences were detected at later time points (Fig. S5A). Similar colonization levels were observed between strains in the urine, bladder, and kidneys at 24 hpi. However, the Δ*katA* mutant exhibited reduced spleen colonization at this time point (Fig. 7A), which could indicate a defect either in dissemination from the urinary tract to the bloodstream or decreased survival once within the bloodstream. Mice from both infection groups exhibited increased bacterial burden and disease severity at 96 hpi compared to 24 hpi, although there were no significant differences between infection groups at this time point (Fig. 7B and Table 1). Thus, while the Δ*katA* mutant exhibited reduced spleen CFUs at 24 hpi, pathogenesis was equivalent to wild-type by 96 hpi.

**Figure 7:**
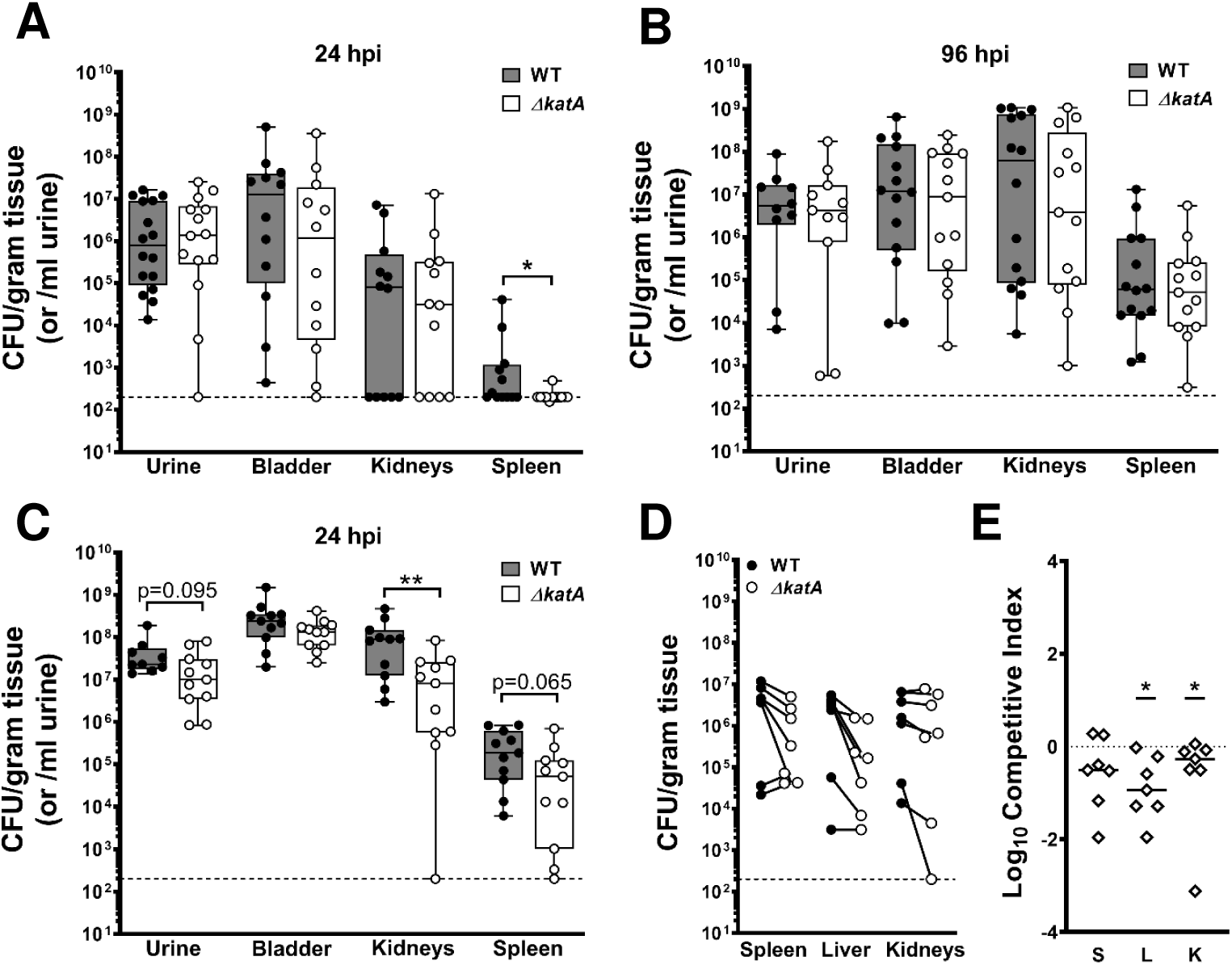
katA contributes to P. mirabilis CAUTI and bacteremia. (A-B) Female CBA/J mice were transurethrally inoculated with 50 µl of a bacterial suspension containing ∼2 x 10^6^ CFU/ml of either WT or Δ*katA*, and a 4-mm segment of silicone catheter tubing was advanced into the bladder during inoculation. Mice were euthanized at (A) 24 and (B) 96 h post inoculation (hpi) for determination of bacterial CFUs in the urine, bladder, kidneys, or spleen. Each black (WT) and white (Δ*katA*) circle represents the log10 CFU per milliliter of urine or gram of tissue from an individual mouse, and the dashed line indicates the limit of detection. * P < 0.05, ** P<0.01 by nonparametric Mann-Whitney test. (C) Female CBA/J mice were transurethrally inoculated via insertion of a 4-mm segment of silicone catheter tubing that had been pre-colonized by either WT or Δ*katA* for 12 h to establish a biofilm (∼2 x 10^6^ CFU/ml). Mice were euthanized at 24 hpi for analysis of CFUs as above. (D-E) Female CBA/J mice were inoculated via tail vein injection of 100 µl of a bacterial suspension containing 1×10^8^ CFU/ml of a 1:1 mixture of wild-type *P. mirabilis* and the Δ*katA* mutant. Mice were euthanized at 24 hpi for determination of bacterial CFUs in the spleen, liver and kidneys. (D) Each black (WT) and white (Δ*katA*) circle connected with a solid line represents the log10 CFU per gram of tissue from an individual mouse. (E) A competitive index (CI) was calculated for the Δ*katA* mutant on a per-mouse basis for the spleen, liver, and kidneys from the ratio of mutant to wild-type recovered from the organ divided by the ratio of mutant to wild-type in the inoculum. Each data point represents the Log10 CI from an individual mouse. Solid lines represent the median, dashed line indicates a competitive index of 1, or a 1:1 ratio of mutant to wild-type. *P < 0.05 by Wilcoxon signed rank test.

**Table 1:** CAUTI disease severity. All mice were assessed for the following parameters at the time of urine collection and euthanasia: bladder hematoma and/or blood in urine, kidney hematoma, kidney color change and/or mottling, and kidney stones. “Any abnormality” indicates whether at least one of these parameters was detected in a mouse. Data represent the number of mice positive for the above parameters out of the total mice infected for each strain during (A) 24h and (B) 96h standard CAUTI model, (C) 24h pre-colonized CAUTI model, and (D) a comparison of mice infected with WT between the two models at 24h. P values were determined by Chi-square test.

| CAUTI Disease Severity | 24h Standard CAUTI<sup>(A)</sup> |  |  | 96h Standard CAUTI<sup>(B)</sup> |  |  | 24h Pre-col CAUTI<sup>(C)</sup> |  |  | 24h Standard CAUTI vs 24h Pre-col CAUTI (WT comparison)<sup>(D)</sup> |  |  |
| --- | --- | --- | --- | --- | --- | --- | --- | --- | --- | --- | --- | --- |
| | WT | $\Delta katA$ | *P* value | WT | $\Delta katA$ | *P* value | WT | $\Delta katA$ | *P* value | CAUTI | Pre-col | *P* value |
| Bladder hematoma and/or blood in urine | 0/12 | 0/12 | - | 0/14 | 0/13 | - | 8/11 | 0/11 | 0.0004 | 0/12 | 8/11 | 0.0003 |
| Kidney hematoma | 0/12 | 0/12 | - | 0/14 | 0/13 | - | 2/11 | 0/11 | 0.1380 | 0/12 | 2/11 | 0.1221 |
| Kidney color change and/or mottling | 0/12 | 0/12 | - | 3/14 | 3/13 | - | 8/11 | 5/11 | 0.1930 | 0/12 | 8/11 | 0.0003 |
| Kidney stone | 0/12 | 0/12 | - | 5/14 | 2/13 | 0.2280 | 4/11 | 0/11 | 0.0270 | 0/12 | 4/11 | 0.0215 |
| Any abnormality | 0/12 | 0/12 | - | 7/14 | 4/13 | 0.3090 | 9/11 | 5/11 | 0.0760 | 0/12 | 9/11 | 0.0001 |

A caveat to the CAUTI model is that the bacterial inoculum is introduced as a suspension, and biofilm formation occurs concurrently with defense against the host immune response. In order to isolate the contribution of the Δ*katA* biofilm defect to *P. mirabilis* pathogenesis during CAUTI, we next inoculated mice with catheter segments that had been pre-colonized for 12 hours by either wild-type *P. mirabilis* or the Δ*katA* mutant to establish catheter biofilms. Importantly, pre-colonizing the catheter segments for 12 hours resulted in a similar inoculum as injection of a bacterial suspension (∼1 x 10^5^ CFU) but caused a more robust infection with pronounced weight loss and disease severity after just 24 hpi (Fig. S5B and Table 1C), such that the experiment could not be extended past this time point. Mice infected with wild-type *P. mirabilis* pre-colonized catheters exhibited more uniform urine and bladder CFUs than in the traditional CAUTI model (Fig. 7C vs 7A), as well as a significantly higher incidence of hematuria and hematoma, kidney discoloration and mottling, and macroscopically-visible kidney stones (Table 1). Infection with the Δ*katA* mutant also displayed greater uniformity in urine and bladder CFUs in the pre-colonized model compared to the traditional CAUTI model; however, in comparison to wild-type *P. mirabilis,* the Δ*katA* mutant exhibited a profound decrease in kidney colonization and a significantly lower incidence of hematuria, bladder hematoma, and kidney stones, as well as a trend towards reduced bacterial burden in the urine and spleen (Fig. 7C and Table 1). Based on these data, wild-type displays greater infection severity and the Δ*katA* mutant displays a more pronounced defect when the infection is seeded from a pre-colonized catheter, which underscores the importance of *P. mirabilis* biofilm development and EPS composition for CAUTI pathogenesis.

While the pre-colonized catheter model clearly demonstrates a pathogenesis contribution for catalase stemming from its role in biofilm development, these studies cannot exclude a role for catalase in defense against ROS. Neutrophils are the first and most abundant immune cells recruited to the bladder in response to CAUTI, and neutrophil oxidative burst represents a robust source of ROS (46–48). Neutrophils appear to be the major factor that limits bacterial growth in the urinary tract during the initial 24 hours post infection (49), and many of the defense strategies used by neutrophils rely on oxidative mechanisms involving ROS (48, 50, 51). The oxidative burst of a single neutrophil has been shown to generate 1–10 μM hydrogen peroxide, and neutrophils in suspension (∼10^6^/ml) are capable of producing steady-state concentrations of ∼10 μM (46). Thus, the decreased kidney and spleen CFUs for the Δ*katA* mutant support an additional role for catalase in protection against host defenses during dissemination and possibly survival within the bloodstream, in which biofilm formation is not likely to play a role.

To investigate biofilm-independent fitness of the Δ*katA* mutant, mice were inoculated via tail vein injection of a 1:1 mixture of wild-type *P. mirabilis* and Δ*katA*. All mice exhibited high bacterial burden in the spleen, liver, and kidneys 24 hpi (Fig. 7D), and the Δ*katA* mutant was significantly outcompeted by wild-type *P. mirabilis* in the liver and kidneys indicating a biofilm-independent fitness defect (Fig. 7E). As neutrophil oxidative burst represents a potent source of hydrogen peroxide during CAUTI, we next investigated the contribution of *P. mirabilis* catalase to defense against opsonophagocytic killing by neutrophils. Interestingly, the Δ*katA* mutant was no more susceptible to opsonophagocytic killing than wild-type *P. mirabilis* (Fig. S6). These data indicate that the contribution of catalase to fitness within the bloodstream is likely due to factors other than defense against neutrophil oxidative burst.

## Discussion

The high degree of resistance against antimicrobials and host defenses that is conferred by bacterial biofilms represents a substantial challenge for the effective treatment of medical device-related infections, including CAUTI. Catheter biofilms containing *P. mirabilis* pose an additional challenge as they are typically comprised of bacterial aggregates, EPS, and crystalline deposits as a result of urease activity, and these crystalline biofilms can result in complete obstruction of urine flow (6). In general, *P. mirabilis* clinical isolates also exhibit a high degree of antimicrobial resistance, including intrinsic tolerance to tetracycline and polymyxin class drugs, acquisition of resistance to aminoglycosides and fluoroquinolones, and the spread of extended-spectrum beta-lactamases and carbapenemases (52–55). Combined with the added resistance provided by the biofilm mode of growth, these properties collectively contribute to the high mortality rate for *P. mirabilis* CAUTI and bacteremia (56–60). Investigation of the processes that contribute to *P. mirabilis* biofilm development and antimicrobial resistance are therefore critical to uncover new potential targets for prevention and treatment.

*P. mirabilis* has long been characterized as having potent catalase activity, and its single catalase gene (*katA*) has been implicated as contributing to infection in three prior genome-wide screens: 1) *katA* expression was upregulated during infection in a mouse model of UTI (42); 2) *katA* was identified as a candidate fitness factor during both single-species and polymicrobial CAUTI (43); and 3) *katA* was detected as an infection-specific fitness factor for survival in the bloodstream (44). We therefore endeavored to delineate the contribution of catalase to several stages of *P. mirabilis* pathogenesis. Here, we show that disruption of catalase results in a slight increase in ROS levels during growth in broth culture, which does not appear to induce oxidative stress in *P. mirabilis* but does increase sensitivity to additional peroxide insult. We also uncovered a critical role for peroxide detoxification via catalase activity in the production of a mature biofilm by *P. mirabilis.* Specifically, loss of catalase activity decreased biofilm biomass due to a severe alteration of EPS composition, particularly the carbohydrate fraction. Importantly, this EPS defect rendered the Δ*katA* biofilms more susceptible to antibiotics and also decreased colonization and dissemination in a mouse model of CAUTI.

The exact sugar moieties that comprise the EPS of *P. mirabilis* strain HI4320 have yet to be fully elucidated; however, hexose moieties previously identified in the LPS outer core of *P. mirabilis* (glucose, galactose, and N-acetyl glucosamine) are likely involved (61, 62). Further exploration is needed to identify the specific carbohydrate moieties that are decreased in the Δ*katA* mutant, and to pinpoint the steps at which loss of catalase activity impacts carbohydrate biosynthesis and EPS production. Regardless, the decrease in EPS-associated carbohydrates clearly resulted in increased antibiotic susceptibility of the Δ*katA* biofilm. EPS defects have been demonstrated to impact biofilm formation and antibiotic susceptibility of other Gram-negative bacteria. For instance, *katA* is important for *P. aeruginosa* biofilm development and biofilm-dependent resistance of treatment with high concentrations of H_2_O_2_ (63), and biofilms formed by *P. aeruginosa Δpsl* (polysaccharide synthesis locus) mutants exhibit increased antibiotic susceptibility (64, 65). Our data also fit within existing literature that have revealed a contribution of catalase to biofilm architecture (66). In *Mycoplasma pneumoniae* (an organism that lacks superoxide dismutase and catalase), treatment with catalase enhanced biofilm formation and altered biofilm structure, such that fewer and smaller tower structures were produced and the resulting biofilms were smoother and more homogenous (3). While treatment of wild-type *P. mirabilis* with bovine catalase did not impact total biofilm biomass, our electron microscopy images suggest that it may increase EPS production and biofilm smoothness. Full determination of *P. mirabilis* HI4320 EPS carbohydrate composition will be necessary to reveal which components are altered when catalase is disrupted, and which (if any) are increased in the wild-type strain upon supplementation with excess catalase.

We also revealed that disruption of *P. mirabilis* catalase results in a biofilm-dependent defect in a mouse model of CAUTI and a biofilm-independent defect in fitness within the bloodstream, thereby demonstrating two separate contributions of the *P. mirabilis* catalase enzyme to pathogenesis. Pre-colonizing the catheter resulted in greater overall bacterial colonization during CAUTI than delivery of a bacterial suspension, which highlights the importance of biofilm formation in CAUTI pathogenesis. The pre-colonized catheter also resulted in greater infection severity overall, as seen by the increased incidence of hematuria and hematoma, kidney discoloration and mottling, and macroscopically-visible kidney stones. These results are in agreement with other investigations concerning the pathogenic potential of biofilm-associated bacteria and bacterial populations that are newly dispersed from a biofilm (66). For instance, *Streptococcus pneumoniae* that have been dispersed from a biofilm have distinct phenotypic properties that differ from those of either biofilm-associated or broth-grown planktonic bacteria, including differential gene expression and increased ability to disseminate and cause infection (67).

The colonization defect of the Δ*katA* mutant that was observed in the spleen during both CAUTI models is also notable, as it could indicate a reduced capacity of the mutant to disseminate from the urinary tract to the spleen, a defect in resistance against host defenses and survival within the bloodstream, or a combination thereof. Since the Δ*katA* mutant also exhibited a fitness defect during direct competition with the wild-type strain in a bloodstream infection, the spleen colonization defect likely pertains to a reduced capacity to survive host immune defenses, such as neutrophil oxidative burst. However, the Δ*katA* mutant was no more susceptible to neutrophil opsonophagocytic killing than the parental strain. One possible explanation for this observation is that neutrophil-mediated killing of *P. mirabilis* may be largely due to non-oxidative mechanisms, such as granule fusion rather than oxidative burst (68–70). Alternatively, *P. mirabilis* catalase may be more critical for resistance of oxidative burst from other cell types, such as macrophages (71, 72). A third possibility is that the slight increase in peroxide levels that occurs in the Δ*katA* mutant may render it more sensitive to other non-oxidative host defenses. Further exploration will be necessary to fully elucidate the role of *P. mirabilis* catalase in immune evasion.

In summary, peroxide detoxification via catalase activity is a critical component of *P. mirabilis* pathogenesis due to a role in biofilm development, antimicrobial resistance, and resistance of host defenses. Elucidation of the underlying mechanism through which catalase impacts carbohydrate and EPS production may uncover new strategies for preventing *P. mirabilis* biofilm formation, disrupting existing biofilms, or increasing sensitivity to ROS and antimicrobial agents. Such strategies could be particularly beneficial for patient populations requiring long-term catheterization.

## Materials and Methods

### Ethics Statement

All animal protocols were approved by the Institutional Animal Care and Use Committee (IACUC) at the State University of New York at Buffalo, Jacobs School of Medicine and Biomedical Sciences (MIC31107Y), and were in accordance with the Office of Laboratory Animal Welfare (OLAW), the United States Department of Agriculture (USDA), and the guidelines specified by the Association for Assessment and Accreditation of Laboratory Animal Care, International (AAALAC, Intl.).

### Bacterial Strains and Culture Conditions

*Proteus mirabilis* strain HI4320 was previously isolated from the urine of a catheterized patient (73). The *P. mirabilis ΔkatA* mutant was constructed in strain HI4320 by the insertion of a kanamycin resistance cassette into the *katA* gene according to the Sigma TargeTron group II intron protocol, as previously described (74). Mutants were verified by selection on kanamycin, PCR, and catalase foam height assay (see below). As insertion of a kanamycin cassette via TargeTron retrohoming can result in polar effects, the Δ*katA* mutant was complemented by PCR-amplifying the wild type *katA* gene and 500 bp of flanking sequences, ligating into a linearized pGEN-MCS vector, electroporation, and selection on ampicillin. Primers used for the generation of the Δ*katA* mutant and Δ*katA+* complement are provided in Supplementary Table 1. *P. mirabilis* HI4320 (wild-type) and the Δ*katA* mutant were routinely cultured at 37°C with aeration in 5 mL of LB broth (10 g/L tryptone, 5 g/L yeast extract, 0.5 g/L NaCl) or on Low salt LB agar (10 g/L tryptone, 5 g/L yeast extract, 0.1 g/L NaCl, 15 g/L agar). Biofilm assays were performed using LB broth (10 g/L tryptone, 5 g/L yeast extract, 0.5 g/L NaCl) or filter-sterilized human urine (Cone Bioproducts, Sequin, TX).

### Catalase Foam Height Assay

100 µl of an overnight culture of each strain was added to a 12 x 75 mm polystyrene tube. Subsequently, 100 µl of 1% Triton X-100 and 100 µL of 30% v/v hydrogen peroxide were added to the tubes, mixed thoroughly, and incubated at room temperature for ∼5 minutes. Following completion of the reaction, the height of O2-forming foam in the test tube was observed.

### Growth Curves

Bacterial overnight cultures were diluted 1:100 into fresh LB and grown in 96-well plates at 37°C with aeration via double orbital shaking in a BioTek Synergy H1. Growth was assessed by measurement of OD_600_ at 15-minute intervals for 18 hours. For hydrogen peroxide sensitivity growth curves, indicated concentrations of hydrogen peroxide were added to the media at the time of inoculation.

### Urease Assay

Urease activity was measured as described previously (45). Briefly, overnight cultures of bacteria were washed once in sterile saline and adjusted to ∼5 × 10^9^ CFU/ml. Cultures were then diluted 1:10 in filter-sterilized human urine supplemented with 0.001% wt/vol phenol red and 500 mM urea and dispensed into the wells of a clear-bottom 96-well plate. Absorbance (OD_562_) was measured every 30 sec for a total of 90 min in a BioTek Synergy H1.

### Motility Assays

Swimming motility agar plates (MOT; 10 g/L tryptone, 0.5 g/L NaCl, 3 g/L agar) were stab inoculated with an overnight culture of wild-type *P. mirabilis* or Δ*katA* mutant and incubated without inverting at 30°C for 16 h prior to the measurement of swimming diameter. Swarming was assessed by inoculating 5 µl of an overnight culture of wild-type *P. mirabilis* or Δ*katA* mutant onto the surface of a swarm plate (LB agar with 5 g/L NaCl), allowing the inoculum to soak in for ∼10 min, and incubating at 37°C for 16 h prior to the measurement of the diameter of each swarm ring.

### ROS Quantification

The ROS-Glo™ H_2_O_2_ Assay Kit (Promega^TM^) was used per the manufacturer’s specifications, with the following modifications to measure the basal level of H_2_O_2_ present in the wild-type and Δ*katA* mutant strain in broth culture. Overnight cultures of each strain were centrifuged, washed once with fresh LB, adjusted to the same optical density, diluted 1:100 in fresh LB, and incubated at 37°C with aeration for 4 hours. After the 4 hour incubation, 80 µl of each sample was added to a 96 well plate with 20 µl of derivatized luciferin substrate and incubated for 30 min at 37°C. To generate a standard curve, a series of Δ*katA* samples were supplemented with increasing concentrations of H_2_O_2_ or an equivalent volume of distilled H_2_0. 100 µl of ROS-Glo™ detection solution was then added to each well, incubated at room temperature for 20 min, and the relative luminescence units (RLU) were read in a BioTek Synergy H1.

### Biofilm Formation for Determination of Viability

Static biofilm formation was performed in tissue culture treated 24-well plates (Falcon 353047) by introducing ∼2 x 10^7^ CFU/ml of either wild-type *P. mirabilis* or the Δ*katA* mutant into 1.5 mL LB broth per well and incubating at 37°C for 20 h. Blank wells for normalization contained only assay medium (LB broth or urine). Following incubation, supernatants were removed and biofilms were washed twice with 1x PBS to remove any remaining planktonic bacteria. A volume of 1.5 mL of sterile PBS was then added to each well and biofilms were scraped with a sterile micropipette tip to resuspend viable bacteria contained within each biofilm. Samples underwent serial 10x dilutions and were spiral plated (Eddy Jet 2, Neutec Group inc, Farmingdale, NY) onto low salt LB agar for enumeration of CFU using a ProtoCOL 3 automated colony counter (Synbiosis).

### Biofilm Biomass Analysis

20 h biofilm formation was performed as above. Where indicated, biofilms were supplemented at time of inoculation with either 0.7 mg of active bovine catalase liver enzyme (∼40,000 units) or 0.1% v/v (∼109 mM) of H_2_O_2_. Supernatants were then carefully removed from wells of a 24-well plate using a VIAFLO Voyager automated pipette (Integra Biosciences, Hudson, NJ), inserted slowly at a 45° angle while making sure to avoid touching the sides and bottom of wells, and set to a speed of 2 for gentle aspiration. Biofilms were air dried for 5 min, stained with 1.5 mL 0.1% crystal violet for 10 min, washed once with distilled H_2_O to remove excess stain, and solubilized in 2 mL of 95% ethanol for 10 min on an orbital shaker at 210 rpm. The wells of the 24-well plate containing the biofilms were then scraped and resuspended with a micropipette tip to ensure all stained biofilm material was solubilized. Crystal violet absorbance (OD_570_) was measured in a BioTek Synergy H1, and blanked using cell-free control wells for each biofilm assay.

### Catheter Biofilm Biomass Analysis

Biofilms were established in 24-well plates as described above, and a 15 mm segment of silicone Foley catheter (Bard 57165814, size 14 French) was placed in the wells at the time of inoculation. After 20 h, catheter segments were transferred into empty wells of a new 24-well plate, all residual LB supernatant was removed from the catheter lumen, and the segments were air dried for 15 min. The segments were then stained with 1.5 mL 0.1% crystal violet for 10 min, washed by gently dunking in distilled H_2_O to remove excess stain, and solubilized in a microfuge tube containing 1 mL of 95% ethanol. The tubes were then transferred to a water bath sonicator for a total of 10 minutes (vortexing every 5 minutes) to ensure removal of the biofilms from the catheter segments. Tubes were then incubated at room temperature at 220 rpm for 10 minutes to ensure full solubilization. The now-clean catheter segments were removed from the solubilized solution, solubilized solutions received a final vortex, and crystal violet absorbance (OD_570_) was measured in a BioTek Synergy H1. Absorbance was blanked using cell-free catheters for each biofilm assay.

### Antibiotic Susceptibility Assay

The following antibiotics were diluted as per manufacturer specifications and made fresh before each experiment: ampicillin sodium salt (Research Products International A40040), ceftriaxone sodium salt (Cayman Chemical 18866), and ciprofloxacin (Tokyo Chemical Industry C2510). For planktonic bacteria, antibiotic susceptibility was determined by assessing growth in increasing concentrations of each antibiotic via OD_600_ as above. Antibiotic susceptibility of biofilms was determined by establishing biofilms as above. After either 4 h or 20 h (as indicated in the text), supernatants were carefully removed and replaced with fresh LB containing increasing concentrations of each antibiotic. Biofilms were then incubated for an additional 20 h, and viable CFUs/mL were determined as above.

### Scanning Electron Microscopy

Biofilms were established on 12mm circular micro cover glass (VWR) within 24-well plates following the same protocol as above. The supernatants were aspirated using a vacuum manifold fitted with a 300 µL tip, inserted slowly at a 45° angle while making sure not to touch the coverslip. Coverslips were gently transferred via sterile forceps to a new/sterile well and biofilms were fixed for 1 h with 2.5% glutaraldehyde in 0.1 M sodium cacodylate buffer containing 0.075% ruthenium red and 0.075 M lysine acetate, pH 7.2. Samples were rinsed three times with 0.2 M sodium cacodylate buffer containing 0.075% ruthenium red (pH 7.2) and then subjected to graded incubations in 30%, 50%, 75%, 95%, and 100% ethanol. Samples were submerged twice in 100% hexamethyldisilazane and air dried. Scanning electron microscopy (SEM) images were captured at the University at Buffalo South Campus Instrument Center using a Hitachi SU-70 microscope equipped with a tilt stage for side angle views.

### EPS Isolation and Analysis

The following protocol was adapted from Bales et al. (75). Biofilms were established in 24-well plates for 20 h as described above and 20 h planktonic samples of each strain were also grown as above. At the 20 h time point, the planktonic samples were normalized by optical density and 1 ml of each was centrifuged, supernatants were removed, cell pellets were resuspended in sterile Milli-Q H_2_O, and the resulting planktonic suspension (PS) was aliquoted and frozen until future use. Supernatants were removed from the biofilm samples, and all biofilm material from the entire 24-well plate was scraped and resuspended into 3 ml sterile Milli-Q H_2_O. A portion of the resulting biofilm suspension (BS) was aliquoted and frozen until future use, and the rest was transferred to sterile microfuge tubes and treated with 6 µl of 37% formaldehyde solution per 1ml to fix the cells and prevent lysis during subsequent extraction steps. The mixture was incubated at room temperature in a chemical hood with gentle shaking (100 rpm) for 1 hour. 400 µl of 1 M NaOH was then added for each 1 ml of biofilm suspension and incubated at room temperature with gentle shaking (100 rpm) for 3 hours to extract EPS. Cell suspensions were then centrifuged (20,000 rcf) for 1 hour at 4°C. The supernatant containing soluble EPS was transferred to a new sterile microfuge tube, while the cellular pellet was resuspended in 1 ml of sterile Milli-Q H_2_O and the resulting cellular fraction (CF) was aliquoted and frozen until future use. The EPS was then filtered through a Spin-X 0.22 µm centrifuge tube filter (Costar 8160) and dialyzed against sterile milli-Q H_2_O (∼300X’s the volume of the sample) using a 3.5K MWCO Slide-A-Lyzer® Dialysis Cassette (Thermo Scientific). Samples were dialyzed for ∼18 h at room temperature with H_2_O changes at 2 and 4 hours. The resulting dialyzed EPS fractions (EF) were aliquoted and frozen until future use.

For EPS component analyses, all isolated fractions (PS, BS, CF, and EF) were thawed and analyzed for characterization of their chemical compositions using commercially available kits. Total carbohydrate was determined by the Dubois phenol-sulfuric acid method (76) adapted to a 96-well plate assay with the Total Carbohydrate Assay Kit (MAK 104, Sigma-Aldrich), using D-glucose as the standard and absorbance read at a wavelength of 490 nm. Total protein content was determined by the Lowry method (77) adapted to a 96-well plate assay with the Microplate BCA Protein Assay Kit (23252, Thermo Scientific), using bovine serum albumin (BSA) as the standard and absorbance read at OD_595_. Total eDNA was determined by the Quant-iT™ PicoGreen^®^ dsDNA Assay Kit (P11496, Invitrogen™) adapted to a 96-well plate, using double stranded lambda DNA as the standard and relative fluorescence units (RFU) read with 470 nm excitation and 525 nm detection. All readings for chemical composition were performed on a Biotek Synergy H1.

### Mouse model of CAUTI

CAUTI studies were carried out as previously described (43). Briefly, the inoculum was prepared by washing overnight cultures of wild-type *P. mirabilis* and the Δ*katA* mutant in PBS, adjusting each to an OD_600_ of 0.2 (∼2 x 10^8^ CFU/ml), and diluting 1:100 to achieve an inoculum of 2 x 10^6^ CFU/ml. Female CBA/J mice aged 6 to 8 weeks (Jackson Laboratory) were anesthetized with a weight-appropriate dose (0.1 ml for a mouse weighing 20 g) of ketamine-xylazine (100 mg/kg ketamine and 10 mg/kg xylazine) by intraperitoneal (i.p.) injection and inoculated transurethrally with 50 µl of the diluted suspension (1 x 10^5^ CFU/mouse). A 4-mm segment of sterile silicone tubing (0.64-mm outside diameter [o.d.], 0.30-mm inside diameter [i.d.]; Braintree Scientific, Inc.) was carefully advanced into the bladder during inoculation and retained for the duration of the study as described previously (45, 78). At both 24 and 96 hours post infection (hpi), urine was collected, the mice were euthanized, and bladders, kidneys, and spleens were harvested into 5-ml Eppendorf tubes containing 1 ml PBS. Tissues were homogenized using a Bullet Blender 5 Gold (Next Advance) and plated using an EddyJet 2 spiral plater (Neutec Group) for determination of CFU using a ProtoCOL 3 automated colony counter (Synbiosis).

### Mouse model of CAUTI using a pre-colonized catheter

Biofilms of wild-type *P. mirabilis* and the Δ*katA* mutant were formed over a 12 hour period on 4-mm segments of sterile silicone tubing (0.64-mm outside diameter [o.d.], 0.30-mm inside diameter [i.d.]; Braintree Scientific, Inc.) and transferred to PBS to remove non-adherent bacteria, resulting in an inoculum of ∼2 x 10^6^ CFU/ml of biofilm-associated bacteria. Female CBA/J mice aged 6 to 8 weeks (Jackson Laboratory) were anesthetized as described above, and pre-colonized catheter segments were then carefully advanced into the bladder. At 24 hpi urine was collected, the mice were euthanized, and bladders, kidneys, and spleens were harvested and processed as above.

### Mouse model of bacteremia

Bacteremia studies were carried out as previously described (79). Briefly, the inoculum was prepared by washing overnight cultures of wild-type *P. mirabilis* and the Δ*katA* mutant in PBS and diluting in PBS to an OD_600_ of 0.1 (1 x 10^8^ CFU/ml). Female CBA/J mice aged 6 to 8 weeks (Jackson Laboratory) were inoculated by tail vein injection of 100 µl (1 x 10^7^ CFU/mouse) of a 1:1 mixture of the *P. mirabilis* HI4320 and Δ*katA* mutant suspensions. Mice were euthanized 24 hpi, and organs were harvested into 5 ml Eppendorf tubes containing PBS (1 ml for spleens and kidneys, 2 ml for livers). Tissues were homogenized as above and plated onto plain LB agar (total CFU) and LB with kanamycin (Δ*katA* CFU) for determination of CFUs. A competitive index (CI) was calculated as follows for all samples in which bacterial burden was above the limit of detection:

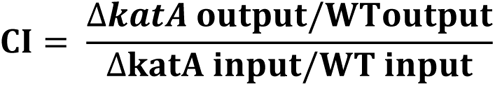

Log_10_CI = 0 indicates that the ratio of the strains in the output is similar to the input, and neither strain had an advantage. Log10CI>0 indicates that Δ*katA* has a competitive advantage over wild-type (WT).

### Statistical Analysis

Normalcy was assessed for all data sets by the Shapiro-Wilk and D’Agostino-Pearson normality tests. Significance was assessed using two-way analysis of variance (ANOVA), nonparametric Mann-Whitney test, unpaired t test, one-sample t test, chi-square test, or Wilcoxon signed rank test, as indicated in the figure legends. All P values are two tailed at a 95% confidence interval. All analyses were performed using Prism, version 7.03 (GraphPad Software, San Diego, CA).

## Acknowledgments

We would like to thank members of the Department of Microbiology & Immunology in the Jacobs School of Medicine and Biomedical Sciences at the University at Buffalo for helpful comments and critiques. This work was supported by the National Institutes of Health [R00 DK105205 and R01 DK123158 to C.E.A.]. The sponsors were not involved in the study design, methods, data collections, analysis, or preparation of the paper. The content is solely the responsibility of the authors and does not necessarily represent the official views of the funders.

